# The influence of kinesthetic motor imagery and effector specificity on the long-latency stretch response

**DOI:** 10.1101/510271

**Authors:** Christopher J. Forgaard, Ian M. Franks, Dana Maslovat, Romeo Chua

**Affiliations:** School of Kinesiology, University of British Columbia; The Brain and Mind Institute, University of Western Ontario; Department of Psychology, University of Western Ontario

## Abstract

The long-latency “reflexive” response (LLR) following an upper-limb mechanical disturbance is generated by neural circuitry shared with voluntary control. This feedback response supports many task-dependent behaviours and permits the expression of goal-directed corrections at latencies shorter than voluntary reaction time. An extensive body of literature has demonstrated that the LLR shows flexibility akin to voluntary control, but it has never been tested whether instruction-dependent LLR changes can also occur in the absence of an overt voluntary response. The present study used kinesthetic motor imagery (Experiment 1) and instructed participants to execute a voluntary response in a non-stretched contralateral muscle (Experiment 2) to explore the relationship between the overt production of a voluntary response and LLR facilitation. Activity in stretched right wrist flexors were compared to standard “not-intervene” and “compensate” conditions. Our findings revealed that on ~40% of imagery and ~50% of contralateral trials, a partial voluntary response “leaked-out” into the stretched right wrist flexor muscle. On these “leaked” trials, the early portion of the LLR (R2) was facilitated and displayed a similar increase to compensate trials. The latter half of the LLR (R3) showed further modulation, mirroring the patterns of voluntary response activity. By contrast, the LLR on “non-leaked” imagery and contralateral trials did not modulate. We suggest that even though a hastened voluntary response cannot account for all instruction-dependent LLR modulation, the overt execution of a voluntary response in the same muscle(s) as the LLR is a pre-requisite for facilitation of this rapid feedback response.

**New and Noteworthy:** We examined volitional modulation of the long-latency stretch response (LLR) using two novel approaches: motor imagery and the execution of contralateral movements. The LLR was only facilitated on imagery or contralateral trials when a voluntary response “leaked-out” into stretched muscle suggesting that a voluntary response in the same muscle as the LLR is a prerequisite for facilitation. Our findings also demonstrate an important distinction between the early (R2) and late (R3) portions of the LLR.

## Introduction

Mechanical perturbations to the arm elicit feedback responses in stretched muscle at short-(SLR: 25-50 ms) and long-latencies (LLR: 50-100 ms) (Hammond, 1956). The SLR is produced by spinal circuitry and is typically only sensitive to background state of the motoneuron pool and features of the perturbation (Pruszynski et al., 2009). By contrast, the LLR receives contributions from both spinal and supraspinal circuitry, providing the motor system with a flexible output given the current state of the limb and the volitional task goal (Pruszynski and Scott, 2012). Among many notable examples, the magnitude of the LLR scales based on target location (Pruszynski et al., 2008), is sensitive to limb mechanics (Kurtzer et al., 2008), and as traditionally investigated, the instruction of how to respond to a perturbation (Hammond, 1956)

The issue of whether volitional characteristics of the LLR are produced through feedback gain changes in presumably supraspinal circuitry (Lee and Tatton, 1975; Pruszynski et al., 2014), or the appearance of a flexible feedback response only arises because a voluntary response is triggered early by the perturbation and superimposes onto the LLR (Rothwell et al., 1980; Manning et al., 2012) has been investigated for many years. Recent studies have consistently shown that LLR modulation begins at a shorter latency than the earliest possible voluntary response (Pruszynski et al., 2008; Forgaard et al., 2015). Despite this, many groups have shown that the magnitude of the LLR remains strongly related to volitional activity (Rothwell et al., 1980; Pruszynski et al., 2008; Manning et al., 2012; Crevecoeur et al., 2013). Scott (2004) has suggested that a distinction between the reflexive and voluntary mechanisms responsible for LLR modulation may be difficult because both responses engage common supraspinal circuitry. The LLR reflects the first volley of activity through the neural circuits that are later engaged by the voluntary response (Scott, 2004; Pruszynski et al., 2008). Thus, the same goal-dependent behaviour observed during the voluntary response is also expressed earlier by the LLR. This could explain the similarities between the LLR and volition and why no satisfactory distinction between this rapid feedback response and voluntary control has been demonstrated.

The purpose of the current study was to determine whether the overt execution of a voluntary response in the same muscle as the LLR is a pre-requisite for instruction-dependent facilitation of this rapid feedback response. In Experiment 1, participants performed kinesthetic motor imagery of a compensate (against the perturbation) task while remaining passive to a wrist extension perturbation. Kinesthetic imagery activates common supraspinal regions as physically executing the imagined response, including areas implicated in the LLR such as M1 (Stinear et al., 2006). In Experiment 2, we investigated whether LLR facilitation requires the specific execution of a voluntary response in the same muscle, or whether facilitation can occur even if a voluntary response occurs elsewhere in the body. Participants were instructed to generate a voluntary response with the left wrist flexors while remaining passive to a right perturbation. We chose this contralateral condition because inter-hemispheric connections between cortical motor regions are known to facilitate the motor pathways leading to homologous musculature of the contralateral limb (Carson et al., 2004). In both experiments, comparisons were made to a passive control condition (DNI) where participants were instructed “do not intervene” and an active condition (ACT) with the instruction to “compensate against the perturbation” (Crago et al., 1976; Manning et al., 2012). We hypothesized that if LLR facilitation could occur in the absence of a voluntary response in the same muscle, the LLR would be enhanced on imagery and contralateral trials (compared to DNI), without any concomitant increases in voluntary activity in right wrist flexors. Alternatively, if LLR facilitation requires the specific execution of a voluntary response in the same muscle, facilitation would be restricted to the trials where a voluntary response occurs in right wrist flexors.

## Methods

### Participants and Ethical Approval

Eighteen (11 female, 7 male, aged 20-33 years) healthy right-handed volunteers participated in this study. The protocol was approved by the University of British Columbia Behavioural Research Ethics Board and conformed to the standards set by the Declaration of Helsinki. Informed written consent was collected before each testing session. After participants signed the consent form, motor imagery abilities were assessed using the Motor Imagery Questionnaire (MIQ-R; Gregg et al., 2010). All participants performed Experiment 1 before Experiment 2; however the order of conditions within each experiment were counterbalanced between participants (practice trials were not counterbalanced; see below). The entire testing session lasted 2 hours and participants were compensated $20 (CAD) for their time.

Due to equipment malfunction during data collection, data were unavailable from one participant. We also found that two participants co-contracted the right wrist flexor and extensor muscles during the (pre-perturbation) baseline epoch of both experiments. These participants had reduced wrist displacement following the perturbation. Thus, analysis for Experiments 1 and 2 focused on data from the remaining 15 participants.

### Experimental Design

After completion of the MIQ-R, participants sat in a height-adjustable chair opposite an oscilloscope placed ~1 meter in front. Both elbows were flexed at 100 degrees and hands were semi-pronated with the wrist joints aligned with the manipulanda rotational axes. Connected to the right manipulandum was a torque motor (Aeroflex TQ 82W-1C). A metal handle adjoined to the motor shaft was placed at the right metacarpophalangeal joints. Padded stops on either side of the wrists were tightened to prevent lateral wrist movement. Custom-molded thermoplastic surrounded the hand and allowed movement around the right wrist while keeping the fingers relaxed. Angular position feedback of the right wrist was continuously provided on the oscilloscope. The home position was 10 degrees of wrist flexion and visually defined on the oscilloscope by arrows. The left manipulandum was immovable and positioned at 10 degrees of wrist flexion (see Manning et al., 2012, for a complete description). Custom molded thermoplastic also surrounded the left wrist and kept the left metacarpophalangeal joints in contact with the manipulandum handle which allowed the monitoring of tension. Prior to commencing practice of the main conditions, 3 maximum voluntary contractions (MVC) of the right and left wrist flexors were collected.

For the practice and testing phases of the experiments, every trial began with an auditory warning signal (80 dB, 50 ms, 500 Hz) and a right wrist extension preload slowly increased (over 500 ms) to 0.25 Nm. Participants were instructed to “resist by lightly activating your right wrist flexors against the load and hold your right wrist at the home position”. Once attaining the home position, participants closed their eyes (Experiment 1 only). A variable foreperiod (2,500-3,500 ms) followed the warning signal and was terminated by a wrist extension perturbation (1.5 Nm, 150 ms). The preload level of extension torque (0.25 Nm) remained for 1000 ms after the perturbation. A minimum of 10 seconds was given between trials.

### Experiment 1

The order of practice conditions for Experiment 1 remained fixed because it was essential that participants understood how to perform the DNI and ACT conditions (and the associated sensations) prior to IMAGERY. All participants began practice with 10 trials of ACT where they were instructed to “compensate against the perturbation as fast as possible”. Participants were encouraged to respond as quickly as possible on these ACT practice trials. This was followed by 10 trials of the DNI condition with the instruction “do not intervene with the perturbation”. Participants then performed 10 trials of IMAGERY where they were instructed “physically do not intervene with the perturbation, but imagine compensating against the perturbation as fast as possible, and the feeling that this would produce”. During practice of these latter two conditions, participants were encouraged to not voluntarily respond with the right wrist. Verbal feedback of “good” was given if no obvious voluntary response (100-200 ms) was present in the wrist flexor EMG or participants were informed that “they responded” when a voluntary response was observed. After the first 30 trials of practice, participants were provided with 2 minutes of rest. Participants repeated the practice sequence a total of 3 times. At the end of practice, all participants reported feeling competent at performing the various conditions.

The testing phase of the experiment was identical to practice trials with the exception of no verbal performance feedback (and counterbalancing of the conditions between participants). During the testing phase, participants performed 10 trials of one condition (e.g., ACT), followed by 10 trials of the next condition (e.g., IMAGERY), and 10 trials of the last condition (e.g., DNI). Two minutes of rest was given prior to beginning the same sequence. Three complete blocks of trials were performed for a total of 90 testing trials. Error trials (e.g., false starts, not beginning at the home target) were recycled online.

### Experiment 2

For Experiment 2, participants were provided with a left wrist target on the oscilloscope that matched the proportion of MVC held during the preload on the right wrist in Experiment 1. This isometric target was a mean of 12.7% (±1.4 standard error of the mean: SEM) MVC. The target was overlaid on the home position target for the right wrist. When the right wrist preload ramped up, participants were instructed to “resist by lightly activating your right wrist flexors against the load and hold your right wrist at the home position. At the same time generate a small contraction in your left wrist flexors such that the left tension line on the oscilloscope also overlays the home position target”.

As in Experiment 1, the order of practice conditions for Experiment 2 remained fixed. All participants began practice with 10 trials of the ACT condition where they were instructed to “compensate against the right perturbation as fast as possible”. Participants were encouraged to respond as quickly as possible. This was followed by 10 trials of DNI practice with the instruction “do not intervene with the right perturbation”. As with Experiment 1, verbal feedback of “good” was provided if no obvious voluntary response (100-200 ms) was present in the wrist flexor EMG or were told that “they responded” when a voluntary response was observed. Participants then performed 10 trials of CONTRA where they were instructed to “physically do not intervene with the right perturbation, but flex your left wrist as fast as possible”. During practice of this latter condition, participants were encouraged to respond as quickly as possible with their left wrist. They were also informed if “they responded on the right” when a voluntary response was observed in right wrist flexors. After the first 30 trials of practice, participants were provided with 2 minutes of rest. Participants repeated the practice sequence 3 times. The testing phase of the experiment was identical to practice trials with the exception of no verbal performance feedback (and counterbalancing conditions). Three complete blocks of testing trials were performed for a total of 90 testing trials. Error trials were recycled online.

### Data Collection and Analysis

EMG data were collected from the muscle bellies of right flexor carpi radialis (FCR), right extensor carpi radialis (ECR), left FCR, and left ECR using pre-amplified surface electrodes connected to an external amplifier (Model DS-80, Delsys Inc., Natick, MA). EMG signals were amplified at 3K and bandpass filtered from 20-450 Hz. Right wrist positional data were collected using a potentiometer (Bourns, Model 6637S-1-103, Riverside, CA) attached to the right wrist manipulandum. Signals were digitized at 2 kHz using a 1401plus data acquisition system and Spike2 (CED) computer software. A customized LabVIEW (National Instruments, Austin, TX) program was used for offline data analysis.

At the beginning of analysis, EMG data were baseline corrected and full-wave rectified. In a manner similar to previous studies (Pruszynski et al., 2008; Ravichandran et al., 2013; Forgaard et al., 2016), right wrist flexor EMG data were divided into 5 epochs relative to perturbation onset. The first epoch occurred from −100 ms until the perturbation onset and was used to obtain a measure of baseline EMG. The second epoch occurred from 25-50 ms post-perturbation onset and captured the SLR (R1 epoch). The third (R2: 50-75 ms) and fourth (R3: 75-100 ms) epochs contained the LLR. The final epoch provided a measure of volitional activity (100-200 ms); however the voluntary response could sometimes begin during the R3 epoch (Ravichandran et al., 2013; Forgaard et al., 2015). Integrated values for each epoch were taken from the rectified EMG traces on a trial-by-trial basis. The mean integrated baseline EMG while holding the 0.25 Nm preload (for each participant) was used to normalize the respective data from the R1, R2, R3, and voluntary epochs. For normalization of the 25 ms duration R1, R2, and R3 epochs, the baseline EMG value (based on 100 ms duration) was divided by 4, whereas no correction was needed for normalization of the 100 ms duration voluntary epoch. A normalized value of 1.0 corresponds to integrated area equivalent to the (duration-corrected) baseline epoch.

In order to objectively determine the presence of any voluntary response “leaks” in the right FCR traces on IMAGERY and CONTRA trials, a .95 confidence interval (CI) was established around the mean of activity for each participant’s normalized voluntary epoch for the DNI condition within each experiment. Any IMAGERY or CONTRA trials with voluntary epoch activity that exceeded the value established by the upper-bound of the .95 CI for DNI were classified as IMAGERY_Leaked_ or CONTRA_Leaked_. It is important to note that often these voluntary response leaks were subtle, and the experimenter did not notice them during data collection (they were smaller than the obvious voluntary responses observed during early practice). Upon inspection of IMAGERY and CONTRA trials during analysis, it was noted that voluntary response leaks were restricted to the 100-200 ms time period in wrist flexors.

For the left wrist flexor data, we were interested in determining premotor RT on CONTRA trials. Left FCR rectified EMG traces were displayed on a computer monitor with a superimposed marker indicating the point where activity increased to more than 3 standard deviations above baseline and remained above this level for greater than 10 ms. Onset latencies were visually verified (and adjusted if necessary) to the onset of the correct muscle burst.

### Statistical Analysis

All statistical analysis was performed using SPSS software. For each predefined response epoch (baseline, R1, R2, R3, voluntary) of right wrist flexor EMG, repeated measures ANOVA tests were used to compare the effect of Condition (Experiment 1: ACT, DNI, IMAGERY_Leaked_, IMAGERY_Non-Leaked_; Experiment 2: ACT, DNI, CONTRA_Leaked_, CONTRA_Non-Leaked_) on the magnitude of the normalized integrated activity (with the exception of baseline epoch where raw integrated activity was used). This analysis was also performed on left wrist flexor baseline epoch activity for Experiment 2. Mauchly’s test was used to test for any violations to the assumption of sphericity, and if necessary, Greenhouse-Geisser *p*-values are reported (along with uncorrected degrees of freedom). Effect sizes were quantified using partial eta squared 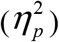 and mean values are reported (± SEM). Dunn-Bonferroni corrected *t*-tests were used as a post hoc test to interpret significant main effects. For Experiment 2, left premotor RT (for CONTRA_Leaked_ and CONTRA_Non-Leaked_) was compared using a paired sample *t*-test. Statistical significance for each test was set at *p* = .05.

## Results

### Experiment 1: Kinesthetic Motor Imagery

The MIQ-R questionnaire revealed that all participants were confident in their capabilities of engaging in various forms of motor imagery. The mean score for internal visual imagery ability was 5.7 (±0.2; between “somewhat easy to see” and “easy to see”) and the mean score for external visual imagery was 5.9 (±0.2; between “somewhat easy to see” and “easy to see”). For kinesthetic imagery ability, the mean score was 5.6 (±0.2; between “somewhat easy to feel” and “easy to feel”). The lowest kinesthetic imagery score was 4.25 (between “neutral” and “somewhat easy to feel”) and the highest score was 7 (“very easy to feel”).

### The influence of kinesthetic motor imagery on feedback responses

The present study was specifically concerned with the relationship between instruction-dependent feedback response modulation and the overt execution of a voluntary response. For Experiment 1, this issue was addressed by instructing participants to engage in kinesthetic imagery of an ACT task while physically not intervening with the mechanical perturbation. This form of mental simulation was ideal for addressing our research question because it is known to produce activation of similar supraspinal circuitry as physical execution of the imagined response (Jeannerod, 2001). We hypothesized that if LLR modulation could occur in the absence of a voluntary response, the LLR would be facilitated on IMAGERY trials, without any concomitant changes to voluntary epoch activity (compared the DNI condition). By contrast, if execution of a voluntary response is required for instruction-dependent LLR modulation, imagery performance would either not influence the LLR, or facilitation would be restricted to trials with a leaked voluntary response.

On average, we observed voluntary response leaks on 39.0% (± 5.7%) of IMAGERY trials (see classification criteria in Methods). One participant did not display any IMAGERY_Leaked_ trials and their data were thus excluded from the subsequent analysis. It is important to note that this participant showed LLR facilitation on ACT trials, and their IMAGERY trials appeared similar to DNI.

Group ensemble displacement profiles are presented in Figure 1A. Group wrist flexor EMG ensemble profiles are presented along a similar time-scale as displacement data in Figure 1B. Enhanced wrist flexor EMG profiles are shown in Figure 2, zoomed-in on the epochs of interest. The *p* values for the Dunn-Bonferroni corrected *t*-test comparisons (following significant omnibus tests) are also presented in Figure 2.

**Figure 1.**
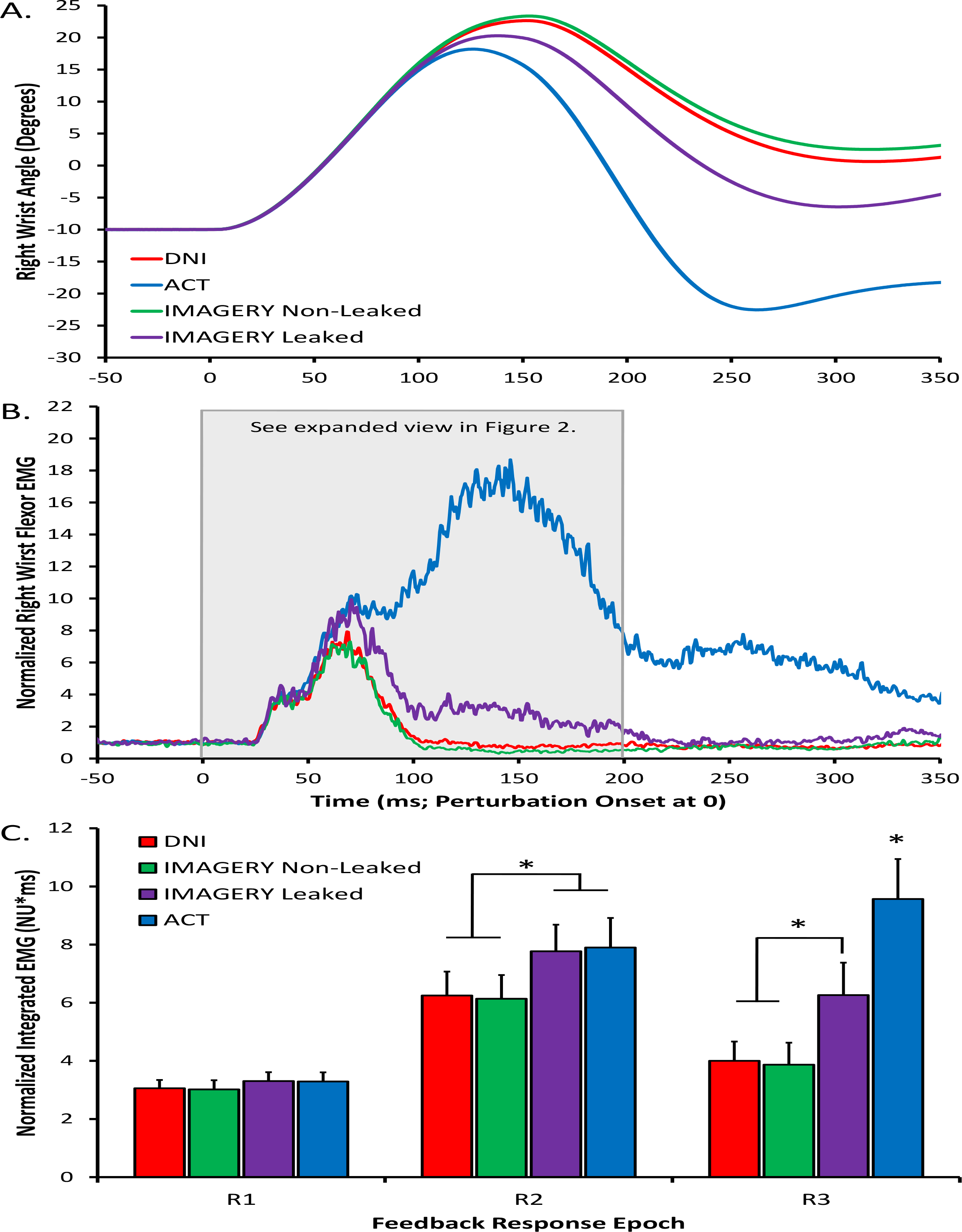
Experiment 1 group right wrist position and right wrist flexor EMG data. DNI: red lines & bars. ACT: blue lines & bars. IMAGERY_Non-Leaked_: green lines & bars. IMAGERY_Leaked_ purple lines & bars. A. Right wrist angle (in degrees). Positive values denote extension and negative values flexion. B. Normalized (to baseline) right wrist flexor EMG along the same time-scale as the displacement data. C. Normalized integrated right wrist flexor EMG for the feedback response epochs of interest. Asterisks denote statistically significant comparisons (*p* < .05). Error bars represent standard error of the mean.

**Figure 2.**
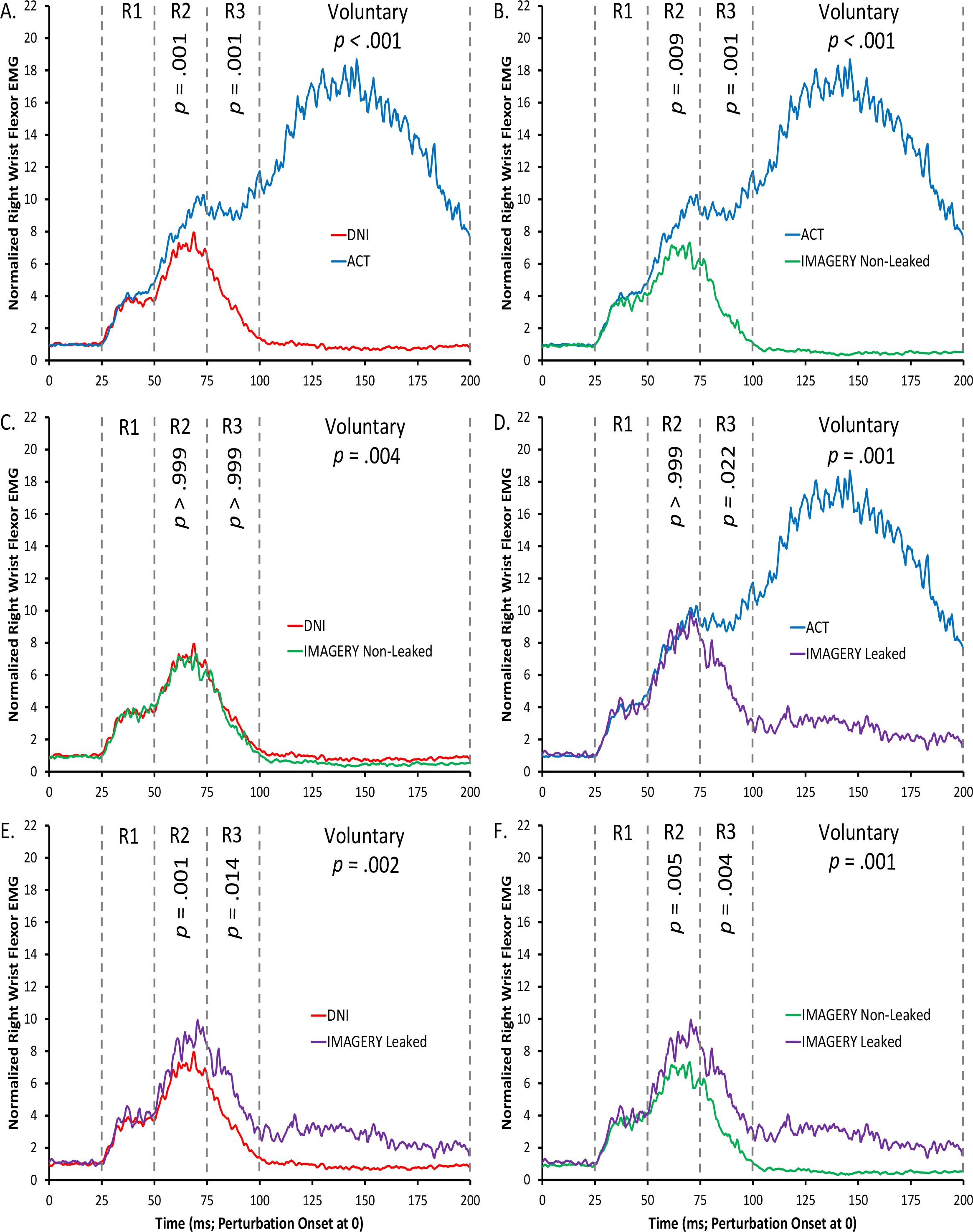
Experiment 1 group ensemble right wrist flexor EMG data, zoomed-in on the epochs of interest. Each panel represents a comparison between different conditions. Following a significant Omnibus ANOVA on a given epoch, Dunn-Bonferroni corrected *p* values are displayed for each comparison.

For studies examining feedback responses following a mechanical stimulus, it is imperative that the background state of the motoneuron pool remains properly controlled prior to perturbation delivery (Pruszynski et al., 2009). An analysis of baseline EMG revealed no significant differences amongst conditions, *F*(3,39) = 1.63, *P* = .198, 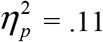. Likewise, no significant differences were seen during the R1 epoch, *F*(3,39) = 2.16, *p* = .108, 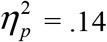. Thus, kinesthetic motor imagery performance did not impact the SLR response (see Figure 1B/C & Figure 2), even on the ~40% of trials where a small voluntary response “leaked-out” into the stretched muscle. This lack of spinal facilitation through mental simulation contrasts the findings of previous motor imagery studies (see Discussion).

Despite no changes to baseline EMG or the SLR response between conditions, motor imagery performance and intended voluntary response execution (ACT) strongly influenced the LLR. An omnibus ANOVA conducted on the R2 epoch revealed the presence of significant differences, *F*(3,39) = 15.91, *p* < .001, 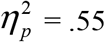. Post hoc analysis showed that activity during this interval was categorically sensitive to the presence of a voluntary response (see Figure 1B/C & Figure 2). For the ACT (7.9 ± 1.1 NU*ms) and IMAGERY_Leaked_ (7.8 ± 0.9 NU*ms) trials, activity was significantly facilitated compared to DNI (6.2 ± 0.8 NU*ms; Figure 2A/E) and IMAGERY_Non-Leaked_ (6.1 ± 0.8 NU*ms; Figure 2B/F). By contrast, DNI and IMAGERY_Non-Leaked_ comparison did not significantly differ (Figure 2C), nor did ACT and IMAGERY_Leaked_ (Figure 2D).

An ANOVA conducted on the R3 epoch also indicated the presence of significant differences, *F*(3,39) = 22.03, *p* < .001, 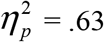. The ACT (9.6 ± 1.4 NU*ms) condition showed the greatest facilitation, followed by IMAGERY_Leaked_ (6.3 ± 1.2 NU*ms; Figure 1C & Figure 2D). IMAGERY_Leaked_ remained significantly larger than DNI (4.0 ± 0.7 NU*ms; Figure 1C & Figure 2E) and IMAGERY_Non-Leaked_ (3.9 ± 0.8 NU*ms; Figure 2F). Critical to our research question, and in line with the R2 findings, DNI and IMAGERY_Non-Leaked_ did not differ significantly (Figure 2C).

During the Voluntary epoch, all conditions differed, *F*(3,39) = 32.49, *p* < .001, 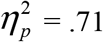. As expected, the largest voluntary response was observed for the ACT condition (14.4 ± 2.4 NU*ms), followed by IMAGERY_Leaked_ (3.2 ± 0.6 NU*ms), DNI (1.1 ± 0.2 NU*ms), and IMAGERY_Non-Leaked_ (0.7 ± 0.2 NU*ms).

To summarize, the findings from Experiment 1 show that volitional intent, including the performance of kinesthetic motor imagery, does not influence the spinally generated SLR. By contrast, the LLR, which receives contributions from multiple supraspinal regions, is strongly influenced by volition. On ACT trials where a large voluntary response always occurred, and the ~40% imagery trials where a small voluntary response “leaked out” into the stretched wrist flexor muscle, a general facilitation was observed during the R2 epoch. The latter portion of the LLR (R3) displayed further modulation on ACT trials and remained significantly facilitated on IMAGERY_Leaked_ trials. On the IMAGERY_Non-Leaked_ trials where a voluntary response was not observed in the EMG recordings, both the R2 and R3 epochs appeared similar to the DNI control condition. These findings strongly suggest that the overt execution of a voluntary response (irrespective of magnitude) is a pre-requisite for instruction-dependent LLR facilitation.

### Experiment 2: Contralateral Responses

#### The influence of effector specificity on LLR facilitation

With the findings of Experiment 1 demonstrating that the execution of a voluntary response was required for LLR facilitation, Experiment 2 sought to determine whether LLR facilitation required the specific execution of a voluntary response in the same muscle (as the LLR), or whether it relied on the (general) execution of a voluntary response anywhere in the body. To test this idea, participants executed voluntary responses with the left wrist flexors while remaining passive to a perturbation applied to the right wrist. During unilateral movements of the upper-limb, inter-hemispheric connections between cortical motor regions facilitate the excitability of motor pathways leading to homologous musculature of the contralateral limb (Carson et al., 2004). It is also known that both the voluntary response and the LLR can be flexibly expressed across limbs during a bimanual coordination task (Mutha and Sainburg, 2009; Omrani et al., 2013). Thus, while we could have addressed our research question by instructing participants to produce voluntary responses in unrelated muscles (e.g., left ankle plantar flexors), this contralateral condition was chosen as it was deemed most likely to result in LLR facilitation in the absence of a voluntary response in stretched muscle.

We hypothesized that if instruction-dependent LLR modulation was related to the general execution of a voluntary response, then the LLR would be facilitated on CONTRA trials, without any changes to voluntary epoch activity in stretched muscle (compared to DNI). By contrast, if instruction-dependent LLR modulation required the specific execution of a voluntary response in the same muscle as the LLR, feedback response facilitation would be restricted to ACT trials, and CONTRA trials where a partial voluntary response “leaked-out” into the stretched muscle.

All 15 participants displayed voluntary EMG leaks in this Experiment. For CONTRA trials, leaked voluntary responses in the right FCR were observed with an incidence rate of 53.2% (±7.2%). As in Experiment 1, the analysis of feedback response data was separated for leaked and non-leaked trials. Group ensemble left wrist tension and right displacement profiles are presented in Figure 3A/B. EMG ensemble profiles are presented along a similar time-scale as tension/displacement data in Figure 3C/D. Right wrist flexor EMG profiles are also shown in Figure 4, zoomed-in on the epochs of interest. Figure 4 also displays Dunn-Bonferroni corrected *p* values following a significant Omnibus ANOVA.

**Figure 3.**
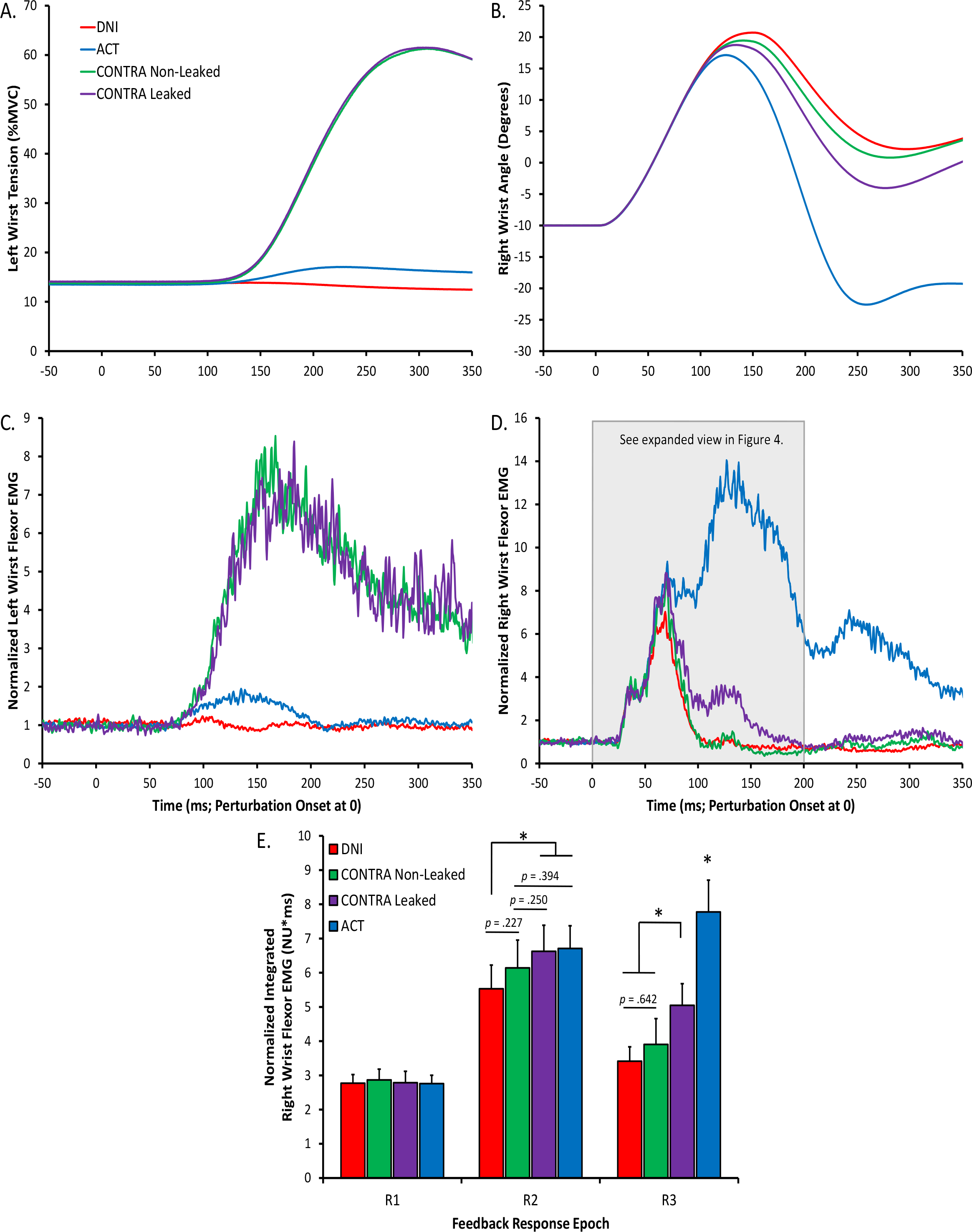
Experiment 2 group data from the left and right wrists. DNI: red lines & bars. ACT: blue lines & bars. CONTRA_Non-Leaked_: green lines & bars. CONTRA_Leaked_ purple lines & bars. A. Left wrist tension (% Maximum Voluntary Contraction). B. Right wrist angle (in degrees). Positive values denote extension and negative values flexion. C. Normalized left wrist flexor EMG ensemble. D. Normalized right wrist flexor EMG ensemble. E. Normalized integrated right wrist flexor EMG for the feedback response epochs of interest. Asterisks denote statistically significant comparisons (*p* < .05). Errors bars represent standard error of the mean.

**Figure 4.**
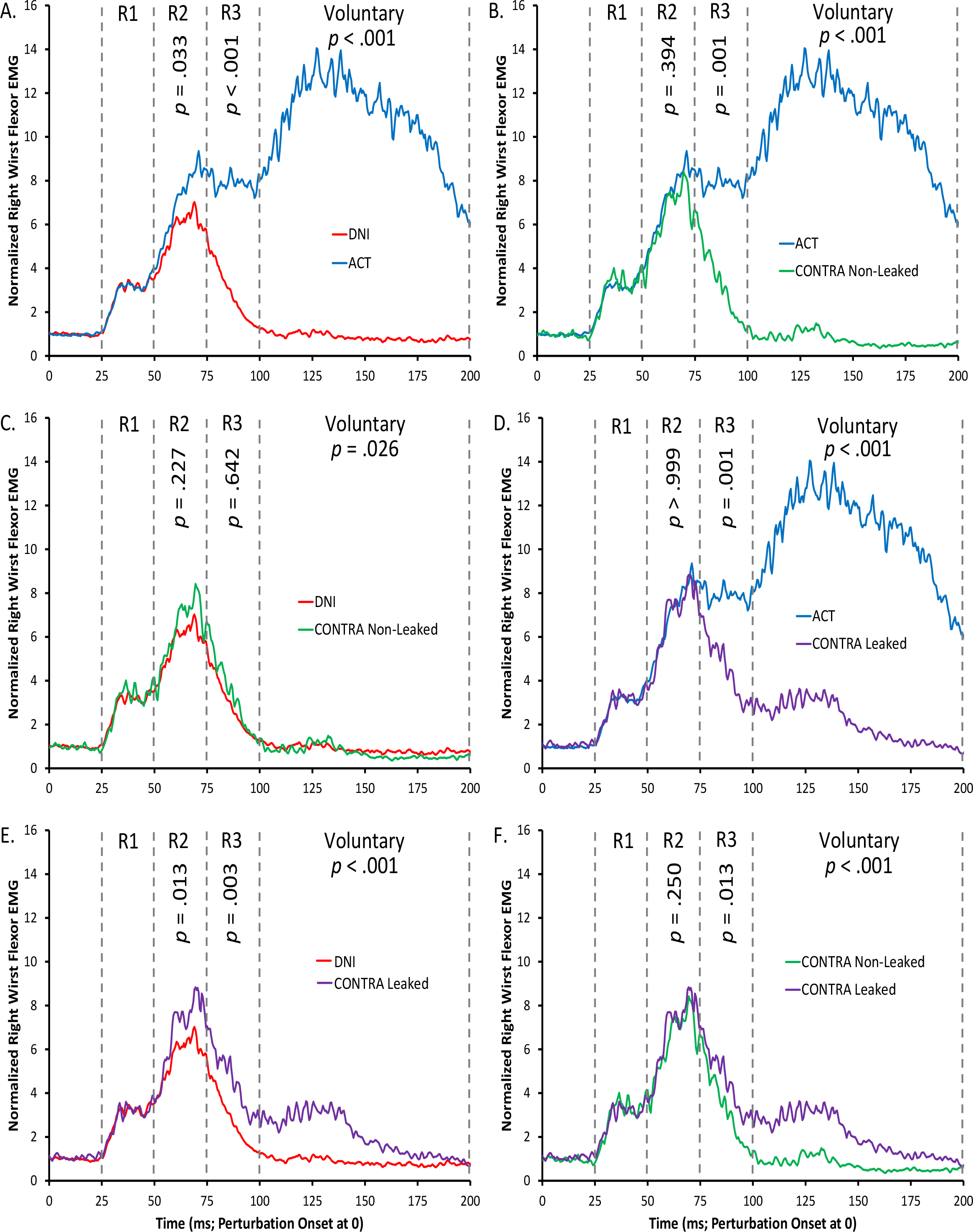
Experiment 2 group ensemble right wrist flexor EMG data, zoomed-in on the epochs of interest. Each panel represents a comparison between different conditions. Following a significant Omnibus ANOVA on a given epoch, Dunn-Bonferroni corrected *p* values are displayed for each comparison.

#### Left wrist flexor activity

Prior to perturbation delivery, participants held an isometric contraction with the left wrist flexors that matched the proportion of MVC with the 0.25 Nm preload on the right wrist. An analysis of left wrist flexor baseline EMG revealed the presence of significant differences, *F*(3,42) = 4.74, p = .006, 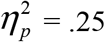. Post hoc analysis showed that activity was increased for the ACT condition (4.3 ± 0.6 mV*ms) compared to CONTRA_Leaked_ (*p* = .010; 3.9 ± 0.6 mV*ms) and CONTRA_Non-Leaked_ (*p* = .025; 3.9 ± 0.6 mV*ms), but did not reach statistical significance when compared with DNI (*p* = .052; 3.9 ± 0.5 mV*ms). Left baseline EMG did not differ (all corrected *p* values > .999) between DNI, CONTRA_Leaked_ and CONTRA_Non-Leaked_. As perturbations were not delivered to the left wrist, we were not concerned with the small baseline increase on ACT trials.

No significant differences were found for premotor RT values for left wrist flexors between CONTRA_Leaked_ (116.2 ± 4.3 ms) and CONTRA_Non-Leaked_ (113.2 ± 5.6 ms) trials, *t*(14) = 0.77, *p* = .450. Although participants were not required to produce a left wrist voluntary response on ACT trials, we did observe leaked responses in left wrist flexors on 44.2% (± 7.9%) of ACT trials. These responses can be seen in the blue profile of Figure 3C. The mean latency of left wrist flexor leaked responses (on ACT trials) was 112.5 ms (± 4.4) ms.

#### Right wrist flexor activity

Despite small baseline differences in the left wrist flexor EMG, no significant differences were found for right wrist flexor baseline activity while participants countered the 0.25 Nm extension preload, *F*(3,42) = 1.50, *p* = .243, 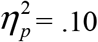. Likewise, no significance differences were found during the R1 epoch, *F*(3,42) = 0.21, *p* = .889, 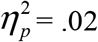 (see Figure 3D/E & Figure 4).

An analysis of the R2 epoch did indicate the presence of significant differences, *F*(3,42) = 7.12, *p* = .001, 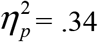. Post hoc analysis showed that similar to Experiment 1, ACT (6.7 ± 0.7 NU*ms) and CONTRA_Leaked_ (6.6 ± 0.8 NU*ms) trials were facilitated compared to DNI (5.5 ± 0.7 NU*ms; Figure 3E & Figure 4A/E). ACT and CONTRA_Leaked_ trials did not differ (Figure 4D). Changes from Experiment 1 were found however, when comparisons were made with the CONTRA_Non-Leaked_ trials (6.1 ± 0.7 NU*ms). This condition was not significantly different from the other three (Figure 4B/C/F).

An ANOVA conducted on the R3 epoch was also statistically significant, *F*(3,42) = 29.03, *p* < .001, 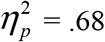, and the pattern of results was identical to Experiment 1. As expected, the ACT condition (7.8 ± 9.6 NU*ms) was larger than all other conditions (see Figure 4A/B/D). The CONTRA_Leaked_ condition (5.0 ± 0.7 NU*ms) was larger than CONTRA_Non-Leaked_ (3.9 ± 0.6 NU*ms; Figure 4F) and DNI (3.4 ± 0.4 NU*ms; Figure 4E). CONTRA_Non-Leaked_ and DNI did not differ significantly (Figure 3E & Figure 4C).

All conditions were significantly different during the Voluntary epoch, *F*(3,42) = 38.93, *p* < .001, 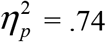 (Figure 4, all panels). The largest voluntary response was observed for the ACT condition (21.9 ± 3.1 NU*ms), followed by CONTRA_Leaked_ (4.8 ± 0.8 NU*ms), DNI (2.2 ± 0.5 NU*ms), and CONTRA_Non-Leaked_ (1.8 ± 0.5 NU*ms).

To summarize, keeping the perturbed limb passive while executing a voluntary response in the contralateral limb does not influence the LLR in stretched muscle. However, during CONTRA trials where a voluntary response leaked out in stretched muscle, the first half of the LLR (R2) was facilitated in a manner similar to ACT trials. These findings suggest that instruction-dependent LLR facilitation is not related to the general execution of a voluntary response (anywhere in the body), rather it is sensitive to the *specific* execution of a voluntary response in the *same muscle* as the LLR.

## Discussion

Our study highlights a strong, indissociable relationship between LLR facilitation and voluntary muscle activation. While it is well established that many features of voluntary control can be expressed during this rapid feedback response (see Pruszynski and Scott, 2012), it has been debated whether the volitional characteristics arise through feedback gain changes or from the superimposition of a voluntary response onto the LLR (e.g., Manning et al., 2012). The current study provides compelling evidence that instruction-dependent facilitation requires the overt execution of a voluntary response specific to the same muscle as the LLR. Across both experiments, the R2 epoch was categorically sensitive to the occurrence of a voluntary response in stretched muscle, whereas activity during R3 mirrored the patterns of activity exhibited by the voluntary response. Here we integrate our findings with other studies and discuss a potential mechanism for instruction-dependent facilitation of this rapid feedback response.

### Imagery and the SLR

With few exceptions (e.g., Wolpaw et al., 1983; Weiler et al., 2018), a majority of evidence has found that the SLR remains insensitive to volition. It is surprising then that previous imagery studies have shown mental simulation can facilitate the SLR (Bonnet et al., 1997; Li et al., 2004; Aoyama and Kaneko, 2011). However, these studies all suffered from the same caveat – not controlling baseline muscle activation prior to the perturbing stimulus, which has a strong influence on the magnitude of the SLR (Pruszynski et al., 2009). It is plausible that the observed spinal modulation by these other groups resulted from subthreshold (or suprathreshold; as reported by Bonnet et al., 1997) changes in motor unit activity prior to the perturbation. Given that background state of the motoneuron pool also influences the LLR (Pruszynski et al., 2009), it was critical that we controlled muscle activity prior to perturbation delivery. Our findings show that when the baseline muscle activity is matched across conditions with a preload, kinesthetic imagery has no influence on the SLR (Figure 2).

### Volition and LLR Facilitation

The main finding from this study is that instruction-dependent LLR facilitation requires the overt appearance of a voluntary response specific to the same muscle. This conclusion is in line with Optimal Feedback Control theory, one tenet of which is that the controller requires an explicit definition of what the motor system is to accomplish (Pruszynski and Scott, 2012). As discussed by Scott (2016), this motor goal is defined by volition or “the self-initiated decision to generate a motor act”. In the context of instruction-dependent modulation, the controller either chooses to remain passive or to compensate against the perturbation. For ACT trials, we believe that selecting the control policy to voluntarily respond modified feedback gains resulting in a facilitated LLR. In our primary conditions of interest (IMAGERY & CONTRA), when a voluntary response leaked out into right wrist flexors (i.e., same musculature as the LLR), we observed a similar R2 facilitation (see Figures 2 & 4). A few previous (non-perturbation) motor imagery studies have also reported leaked responses (e.g., Guillot et al., 2007; Maslovat et al., 2013), but it remains unresolved whether the “leaks” are caused by a failure to completely inhibit the imagined response, or by subliminal activation of motor areas (during imagery) that occasionally produces the overt activation of muscle. While our findings cannot distinguish between these alternatives, the generic R2 increase on all trials with a voluntary response suggests that a similar control policy (to voluntarily respond) was also implemented on leaked trials. Critical to our research question, on the IMAGERY/CONTRA_Non-Leaked_ trials, where no voluntary response occurred in stretched muscle, our findings suggest the feedback gains contributing to the LLR were similar to DNI.

While all conditions with a voluntary response in right wrist flexors showed a similar R2 facilitation, different findings emerged in R3. Other studies have also reported differences between the two phases of the LLR. For example, perturbations delivered during a reaching task produce modulation beginning in R2 (Nashed et al., 2012), but if the participant selects to move to a new target location, modulation is delayed until the R3 epoch (Nashed et al., 2014). In a compensate condition, Lee and Tatton (1975) reported that ~40% of participants displayed two reflexive responses in wrist flexors during the LLR, corresponding closely to R2 and R3. The authors suggested that these were produced by different pathways, the former driven by a transcortical route involving S1 and M1, and the latter produced by a cerebellar-M1 pathway. The R3 epoch can also be influenced by voluntary response superimposition (Ravichandran et al., 2013; Forgaard et al., 2015). Thus, it is plausible in our study that R3 activity mirrored the patterns exhibited during the voluntary epoch due in fact to superimposition of the voluntary response. Although our current methodology cannot distinguish the neural substrates mediating R2/R3, our findings provide evidence that feedback gains can be independently modified during these epochs.

Whereas the SLR is driven by spinal circuitry, multiple spinal and supraspinal sources contribute to the LLR (Lourenço et al., 2006). Many studies have focused on M1 which does contribute to some forms of LLR modulation including the integration of intersegmental limb dynamics (Pruszynski et al., 2011), target-dependent modulation (Pruszynski et al., 2014), environmental stability regulation (Shemmell et al., 2009), and inhibiting the LLR when participants are instructed to “let-go” (Spieser et al., 2010). In contrast, multiple lines of evidence suggest that M1 does not have a role in the LLR facilitation between a true passive (DNI) and a compensate condition. For instance, cortical potentials localized over M1 do not change (based on these instructions; MacKinnon et al., 2000), a TMS-induced silent period of M1 has no influence on LLR facilitation (Shemmell et al., 2009), and intra-cortical inhibition within M1 is similar between instructions (Lewis et al., 2006). The authors of these latter three papers suggested that instruction-dependent LLR facilitation (between compensate and DNI) occurs downstream of M1.

One potential downstream contributor is the reticular formation (Ravichandran et al., 2013), however this brainstem structure is unlikely involved in facilitating the LLR in conditions similar to the current work (Forgaard et al., 2016). The dentate nucleus in the cerebellum (via connections with M1) may play a role, but likely not until R3 (Strick, 1983). A recently implicated candidate is area A5 in the parietal cortex (Omrani et al., 2016). These authors developed a non-human primate behavioural analog to the DNI instruction and compared a compensate task where the animals quickly returned their arm to the initial position. A similar initial response in M1 was observed between conditions (see also Omrani et al., 2014). However, a remarkable finding was that A5 neurons were highly sensitive to task engagement. Compared to the passive condition, on trials where the animals generated corrective motor responses, A5 neurons immediately (~25 ms) increased their firing rate. Changes in other prefrontal and parietal areas did not occur until at least 40 ms after onset of the perturbation, too late to influence the R2 epoch. Omrani and colleagues proposed that A5 was involved in selecting the task engagement control policy.

A5 contributes to movement control via intra-cortical connections with other regions such as M1 and though direct corticospinal projections onto spinal interneurons (Rathelot et al., 2017). Given previous evidence suggesting that instruction-dependent LLR facilitation occurs downstream of M1 (MacKinnon et al., 2000; Lewis et al., 2006; Shemmell et al., 2009), it is plausible that corticospinal inputs from A5 could have acted at a spinal level to cause the generic R2 facilitation observed in our study. For example, spinal mechanisms such as reverberating Ia afferents are known to contribute to the LLR (Hagbarth et al., 1981) and descending inputs could modify this mechanism by various means such as reducing presynaptic Ia inhibition. It is also known that spinal excitability, as tested through the H-reflex, increases ~25 ms prior to the onset of a voluntary response (MacKinnon and Rothwell, 2000). Given that the fastest voluntary RTs in a compensate task occur during the R3 epoch (Forgaard et al., 2018), the generic R2 increase we have observed on trials with a voluntary response may have resulted from descending modulation of spinal excitability preceding the voluntary response. This mechanism could explain why the R1 response can be facilitated when the voluntary response is temporally cued to occur earlier (Lewis et al., 2006), and why the LLR does not modulate if onset of the voluntary response occurs far beyond the end of this stretch response (Manning et al., 2012).

### Concluding Remarks

In this study, we examined instruction-dependent LLR modulation; specifically the facilitation that occurs when participants transition from a true passive condition to one involving active compensation against the perturbation. It has been extensively debated whether this form of LLR facilitation is produced by modifications to feedback gains or from superimposition of a voluntary response. Our study confirms that LLR facilitation occurs earlier than the shortest latency voluntary response but critically, we demonstrate that the overt generation of a voluntary response specific to the same muscle as the LLR remains a pre-requisite for feedback gain changes to occur.

## Acknowledgements

This work was supported by the Natural Sciences and Engineering Research Council of Canada.

